# The contribution of parent-to-offspring transmission of telomeres to the heritability of telomere length in humans

**DOI:** 10.1101/276030

**Authors:** Dayana A. Delgado, Chenan Zhang, Kathryn Demanelis, Lin S. Chen, Jianjun Gao, Shantanu Roy, Justin Shinkle, Mekala Sabarinathan, Maria Argos, Lin Tong, Alauddin Ahmed, Tariqul Islam, Muhammad Rakibuz-Zaman, Golam Sarwar, Hasan Shahriar, Mahfuzar Rahman, Muhammad Yunus, Jennifer A. Doherty, Farzana Jasmine, Muhammad G. Kibriya, Habibul Ahsan, Brandon L. Pierce

## Abstract

Leukocyte telomere length (LTL) is a heritable trait with two potential sources of heritability (h^2^): inherited variation in non-telomeric regions (e.g., SNPs that influence telomere maintenance) and variability in the lengths of telomeres in gametes that produce offspring zygotes (i.e., “direct” inheritance). Prior studies of LTL h^2^ have not attempted to disentangle these two sources. Here, we use a novel approach for detecting the direct inheritance of telomeres by studying the association between identity-by-descent (IBD) sharing at chromosome ends and phenotypic similarity in LTL. We measured genome-wide SNPs and LTL for a sample of 5,069 Bangladeshi adults with substantial relatedness. For each of the 7,254 relative pairs identified, we used SNPs near the telomeres to estimate the number of chromosome ends shared IBD, a proxy for the number of telomeres shared IBD (T_shared_). We then estimated the association between T_shared_ and the squared pairwise difference in LTL ((ΔLTL)^2^) within various classes of relatives (siblings, avuncular, cousins, and distant), adjusting for overall genetic relatedness (ϕ). The association between T_shared_ and (ΔLTL)^2^ was inverse among all relative pair types. In a meta-analysis including all relative pairs (ϕ >0.05), the association between T_shared_ and (ΔLTL)^2^ (*P*=0.002) was stronger than the association between ϕ and (ΔLTL)^2^ (*P*=0.45). Our results provide strong evidence that telomere length (TL) in parental germ cells impacts TL in offspring cells and contributes to LTL h^2^ despite telomere “reprogramming” during embryonic development. Applying our method to larger studies will enable robust estimation of LTL h^2^ attributable to direction transmission.

## INTRODUCTION

Telomeres are DNA-protein complexes at the end of mammalian chromosomes that maintain genome stability by preventing recombination, end-to-end fusion, and DNA damage at chromosome ends ^1^. In somatic cells, shortening of the DNA component of the telomere occurs with each round of cell division due to the “end-replication problem”. In stem cells, extension of telomeres by telomerase counters this shortening, but gradual shortening still occurs with age, with critically short telomeres contributing to cellular senescence. Consequently, leukocyte telomere length (LTL) is considered an indicator of biological aging and a potential biomarker of risk for age-related diseases ^2; 3^. Short LTL is associated with increased risk for several age-related diseases in observational studies, including cardiovascular disease, hypertension, liver disorders, diabetes, atherosclerosis, and overall mortality ^2; 4-7^. In contrast, Mendelian randomization studies suggest that longer telomeres increase risk for several types of cancer, including lung adenocarcinoma ^8^, melanoma ^9^, glioma ^10^, neuroblastoma ^11^, and chronic lymphocytic leukemia ^12^.

LTL is a highly heritable trait ^13-16^ and is known to be effected by inherited variation in non-telomeric regions, e.g., SNPs (single nucleotide polymorphisms) that influence telomere maintenance ^17^. However, LTL has a unique additional potential source of heritability (h^2^): variability in the lengths of telomeres themselves in the parental gametes that produce the offspring zygotes ^18^, under the assumption that gamete telomere length (TL) partially determines TL in adult tissues (including leukocytes and germ cells). This second source of h^2^ is described as “direct” transmission of telomeres (**Figure 1**) and is hypothesized to be the primary source of TL h^2,18^. Telomeres undergo “reprogramming” in the embryo, initially by recombination-based mechanisms and then by telomerase at the blastocyst stage and later ^19^, but is unclear to what extent reprogramming alters the impact of parental germ cell TL on TL in offspring cells.

**Figure 1.**
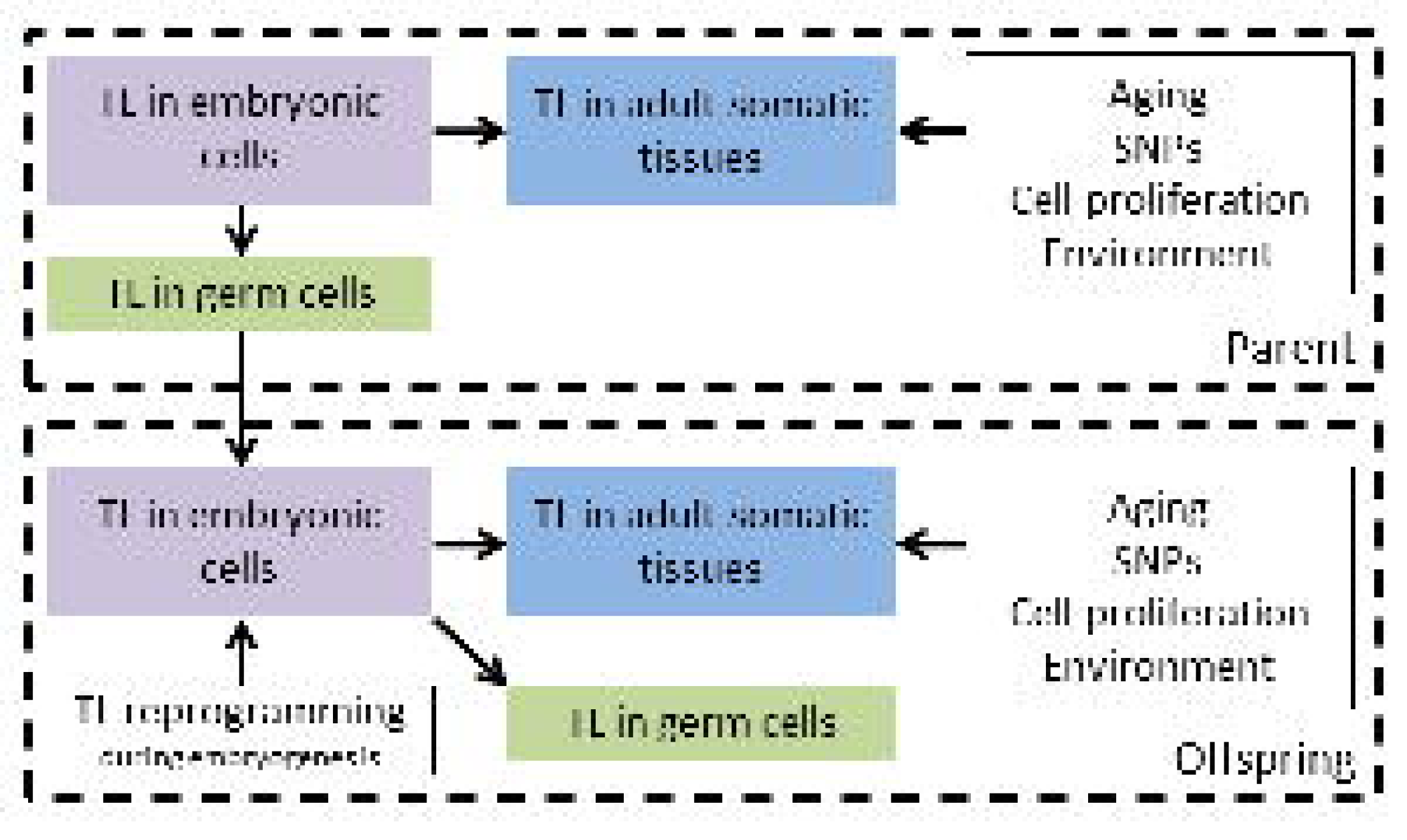
Causal diagram of trans-generational (i.e., “direct”) inheritance of telomere length. TL in adult tissues, including germ cells, is influenced by TL in embryonic cells. TL in embryonic cells of offspring is influenced by the TL in parental germ cells. TL in offspring undergoes “reprogramming” during the pre-implantation stage of embryonic development. Other sources of variation in TL include age, SNPs, cell proliferation rates, and environmental factors.

Evidence supporting an impact of parental TL on offspring TL includes the observed associations of advanced paternal age at conception and increased offspring LTL ^16; 20-26^. This association is often attributed to age-related telomere lengthening in spermatogonia, which express high levels of telomerase ^20^, providing evidence that acquired variation in TL in the male germ line contributes to offspring TL. Also supporting this notion are studies of families with genetic disorders affecting telomere maintenance, in which affected parents who carry a mutation that renders low telomerase activity and short telomeres have non-carrier children whose telomeres appear shorter as compared to the general population, despite not inheriting the mutation ^27-29^. These findings are also supported by data from telomerase deficient mice ^29^.

In light of the evidence supporting the contribution of “direct” transmission of telomeres to LTL h^2^, in this work, we attempt to detect the effect of direct transmission on LTL in a population-based setting. Our approach is to estimate the association between identity-by-descent (IBD) sharing at chromosome ends (a proxy for sharing telomeres IBD) and phenotypic similarity in LTL between relative pairs. Prior family-based and SNP-based h^2^ studies of TL have not attempted to disentangle the effects of “direct” inheritance of TL from other sources of h^2^, and it remains unclear whether the h^2^ of TL is primarily driven by inherited genetic variation in non-telomeric regions (e.g., SNPs that impact telomere maintenance) or whether TL is substantially influenced by the length of the telomeres present in the parental germ cells ^18^. The latter hypothesis implies that TL is not completely “reset” during embryogenesis and early development.

## EXPERIMENTAL METHODS

### Study participants

DNA samples were obtained from participants in the following Bangladeshi studies: The Health Effects of Arsenic Longitudinal Study (HEALS), an expansion of the HEALS cohort (ACE), and The Bangladesh Vitamin E and Selenium Trial (BEST). HEALS is a prospective study of the health outcomes associated with arsenic exposure through drinking local well water in 11,746 adults (18-75 years of age). An expansion of the HEALS cohort (ACE) occurred between 2006 and 2008, and an additional 8,287 participants were recruited. Details of HEALS have been described previously ^30^. BEST is a chemoprevention trial that assessed the effects of vitamin E and selenium supplementation on non-melanoma skin cancer risk. Details of the BEST study have been described previously ^31^. The HEALS and BEST study participants included in this work are primarily from the Araihazar area, a rural community east of Dhaka. Multiple members of extended families were often recruited, resulting in a substantial number of relative pairs in these cohorts. These studies have been approved by the Ethical Review Committee of International Center for Diarrheal Disease Research, Bangladesh, The Bangladesh medical Research Council, and the Institutional Review Boards of the University of Chicago and Columbia University.

### DNA extraction and genotyping

Details of sample collection, DNA extraction and genotyping have been described by Pierce, et al. ^32; 33^. In brief, genomic DNA for HEALS (including ACE) and BEST samples were extracted from clotted blood using the Flexigene DNA Kit (Cat # 51204) and whole blood using the QIAamp 96 DNA Blood Kit (Cat # 51161), respectively. DNA samples were processed on HumanCytoSNP-12 v2.1 chips. Prior to quality control, our genotype data consisted of 5,499 individuals with 299,140 SNPs measured. We removed individuals with call rates < 97% (n = 13), gender mismatches (n =79), and technical replicate samples or duplicates (n = 53). We excluded SNPs that had low call rates, i.e. (< 95%) (n = 20), were monomorphic (n = 39,798), did not have rs identifiers (n = 941), and had HWE p-values < 10^−10^ (n = 634). The HWE threshold used is not stringent because our sample includes relatives, leading to inflated HWE test statistics. In an unrelated set of participants, HWE testing confirmed no SNPs used in the analysis had an HWE P value < 10^−7^.

### Measurement of LTL using qPCR

For 2,203 genotyped HEALS samples, LTL was measured using quantitative(q)-PCR using two different plate designs (Design 1 and 2) on 96-well plates. The relative telomere length is represented as a ratio of the telomere repeat copy number to single-gene (*RPLP0*) copy number, i.e. T/S ratio. Details of this methodology have been described previously ^34^. Plate design 1was based on Ehrlenbach et al. ^35^ and consisted of triplicates of 14 participant DNA samples per plate and six replicates of reference DNA sample, with telomere sequence and *RPLP0* measured on the same plate. Plate design 2 was based on Cawthon et al. ^36^, and we used paired plates for amplification of telomere and single-copy gene of each DNA sample, each consisting of triplicates of 31 participant DNA samples and triplicates of reference DNA sample.

The coefficients of variation (CV) were calculated as the standard deviation of the sample replicates divided by the mean of the replicates. To assess the reproducibility of the measured T/S ratios for Plate Design 1 and 2, we re-ran 37 and 31 samples on separate days and calculated the inter-plate CVs to be 11.7% and 9.8%, respectively.

### Measurement of LTL using QuantiGene Chemistry on a Luminex Platform

The Affymetric-Panomics QuantiGene Plex is a probe-based non-PCR assay developed in our lab to quantify LTL in 1,825 BEST and 1,047 ACE participants. The details of this methodology have been described previously ^37^. The Luminex method has been validated in a blinded comparison to Southern Blot ^38^, the gold standard for TL measurement in DNA samples. Like qPCR, the Luminex-method method produces a relative telomere length measure represented by a ratio of telomere probe to reference gene probe, known as the Telomere Quantity Index (TQI).

### LTL Standardization

To remove experimental variation in our measures of LTL, we utilized linear mixed-effects models (lme package in R), where position and plate were treated as fixed and random effects, respectively. We performed three separate mixed effects models; one for the Luminex-based LTL measures and one for each qPCR plate design. To harmonize the LTL measures from the two experimental methods, we standardized the residuals from mixed-effects regressions and adjusted for age, sex, and cohort in a separate regression model. These standardized residuals were used in all downstream analyses. Additionally, we generated two variables that represent the difference in LTL between relative pairs: the squared difference (ΔLTL)^2^ was our primary outcome (due to its utility for h^2^ estimation (see below)), and the absolute difference |ΔLTL| was our secondary outcome (presented as supplementary results).

### Estimating telomeres shared

We performed LD pruning on 257,747 genotyped SNPs to produce a subset of uncorrelated markers (i.e. linkage equilibrium). In PLINK, we used the “indep-pairwise” command, where we specified three parameters: a 100kb window size, a 25kb count to shift the window at the end of each step, and a pairwise squared correlation (r^2^) threshold of 0.1 to eliminate SNP pairs with an r^2^ > 0.1 in each window, resulting in 52,066 uncorrelated SNPs. Using this set of uncorrelated SNPs, we restricted to the 180 SNPs closest to the end of each chromosome to estimate identity-by-descent (IBD) probabilities for each chromosome end, for all possible pairs of individuals (~12.8 million pairs). We chose to use 180 SNPs because this number gave us very clear separation among pairs who shared exactly 2 chromosome ends, exactly 1 end, and <1 end for all chromosome arms (**Figure S1**). When using fewer than 180 SNPs (e.g., 150 SNPs), for some chromosomes, we did not see a clear gap in the IBD proportion between the pairs sharing exactly one end and the pairs sharing <1 end. The genetic and physical distances spanned by these SNPs at each chromosome end are described in **Table S1** and the frequency of pairs sharing chromosome ends IBD are described in **Table S2**.

We used the IBD proportion for each chromosome end, obtained from PLINK, to define a discrete value for the number of chromosome ends shared IBD, which represents likely sharing of telomeres IBD. For each chromosome end, we defined 0 shared telomeres if the IBD proportion was <0.45, 1 shared telomere if the proportion was between 0.45 and 0.9, and 2 shared telomeres if the proportion was ≥0.9. We summed the number of shared telomeres across all autosomal chromosome p- and q-arms to obtain a total number of ends shared IBD for each participant pair, which we take as an estimate of the number of telomeres shared IBD (T_shared_) (**Figure 2**). We excluded the p-arm of chromosomes 13-15, 21, and 22 from our T_shared_ count, because those arms had a >14 Mb gap between the most telomeric SNP and the chromosome end (as defined by the Genome Reference Consortium Human build 37 (GRCh37/hg19)). For all other chromosome ends, this gap was < 0.3 Mb, except 1p which had a gap of 0.75 Mb. Recombination events are extremely rare in intervals of this size, implying that sharing a segment IBD at a chromosome end (based on 180 SNPs) is a strong proxy for sharing a telomere IBD. Thus, using only 39 chromosome ends results in a theoretical maximum of T_shared_ = 78 (for twins). We divided T_shared_ by 78 to represent the estimated proportion of telomeres shared IBD.

**Figure 2.**
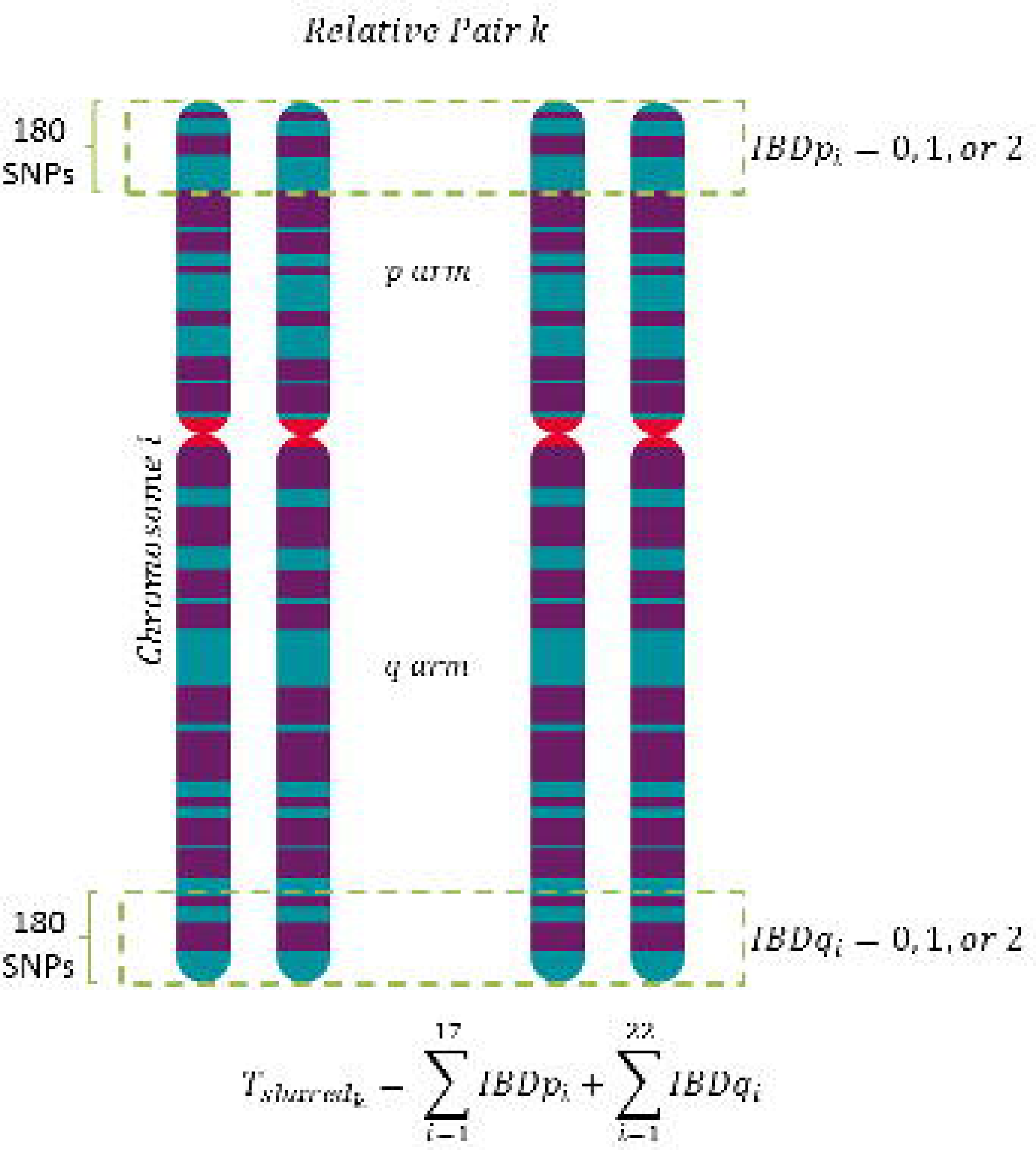
Calculation of T_shared_ for a relative pair. Chromosome end sharing is used as a proxy for telomere sharing. 180 SNPs from each chromosome end were used to calculate the number of ends shared IBD (0, 1, or 2) at each chromosome end for each relative pair. The sum of the IBD count (0, 1, or 2) over all chromosome ends (17 ends p-arm ends (5 excluded) and 22 q-arm ends) resulted in a pair’s T_shared_ value.

We classified each participant pair according to their relationship type (i.e., parent-offspring, sibling, cousin, avuncular, etc.) using the estimated genetic relationship (ϕ) and the IBD probabilities (P(IBD)=0 and P(IBD=1)), all estimated using the PLINK --genome command applied to genome-wide SNPs (**Figure 3**). Paris with ϕ=0.25 could in theory be grandparent-grandchild pairs or half siblings, but based on what we know regarding the demography of the population groups we are studying and the age difference of these pairs, the vast majority of these pairs should be cousins, so we classified them as such. The ϕ threshold separating cousin pairs from “distant” pairs (ϕ=0.125) was somewhat arbitrary due to lack of clear separation between these groups (in terms ϕ) at lower ϕ values; thus, there is likely contamination of the “distant” group with cousin pairs. We conduct sensitivity analyses (see below) to ensure our results are not sensitive to arbitrary decisions regarding this threshold.

**Figure 3.**
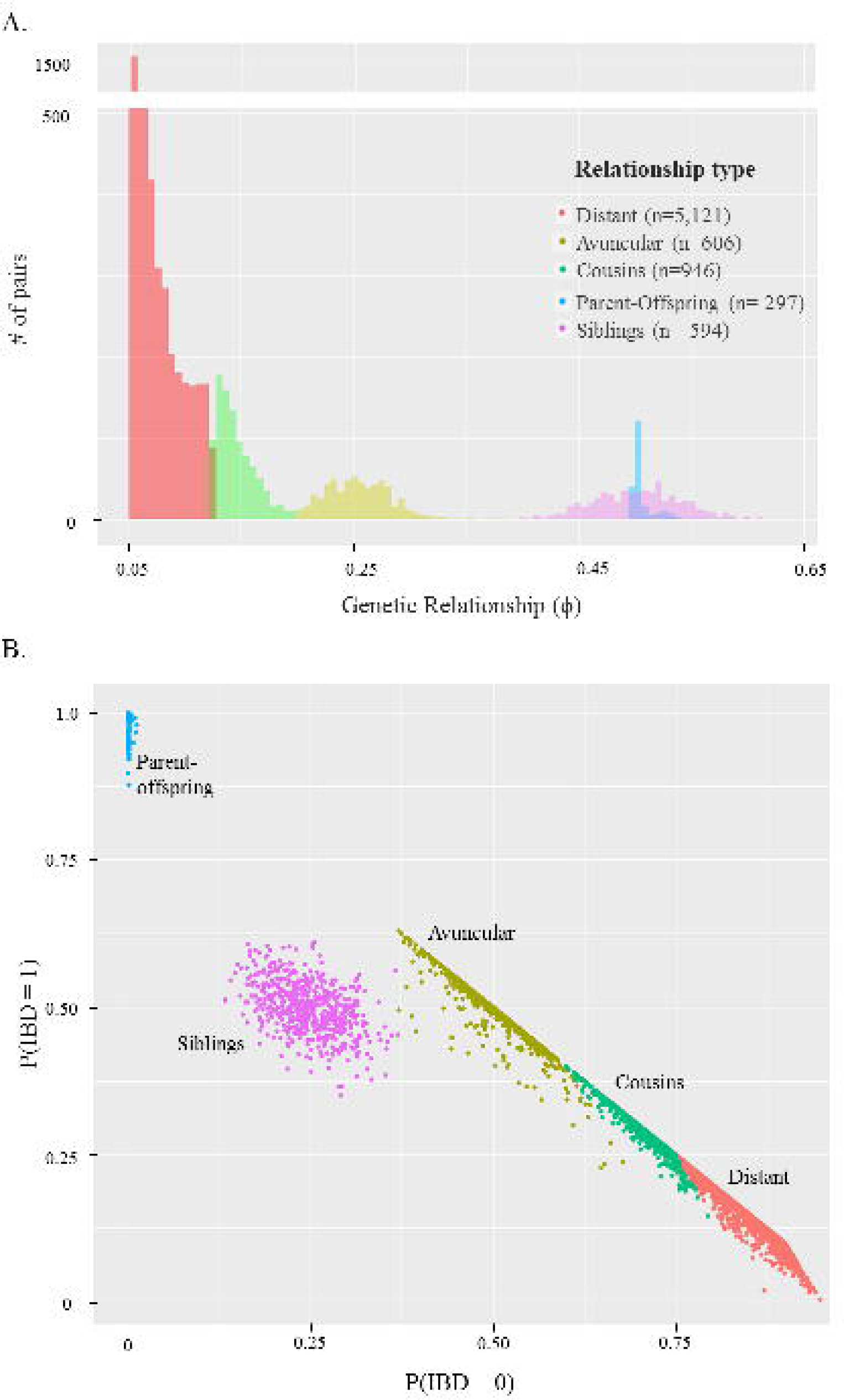
Genetic relationships present in the Bangladeshi cohort. (a) Distribution of the genetic relationship (ϕ) estimated using genome-wide SNPs, restricting to ϕ > 0.05. (b) Scatterplot of P(IBD=0) (x-axis) versus P(IBD=1) (y-axis).

### Statistical Analyses

For each relative pair type, we used linear regression to estimate the association between the T_shared_ proportion and the squared difference in LTL ((ΔLTL)^2^), adjusting for each pair’s ϕ value. This regression was performed for 583 sibling pairs, 605 avuncular pairs, 945 cousin pairs, and 5,121 distant relative pairs. These are quasi-independent pairs, meaning that a given individual can be included in multiple pairs, but pairs are treated as independent. While quasi-independent pairs are routinely used in studies using sib-based IBD methods for h^2^ estimation ^39-41^, we also conducted analyses restricting to pairs with no duplicate individuals. Parent-offspring pairs were not analyzed because we expect no variation in the T_shared_ proportion (or ϕ) among these pairs.

To estimate the association between T_shared_ proportion and (ΔLTL)^2^ using data on all types of relative pairs, we combined the association estimates for each relationship type in a fixed-effects meta-analysis using the ‘metafor’ package in R. Fixed-effects meta-analysis requires two key assumptions: 1) homogenous effect-size across all groups and 2) no between-group variance. Additionally, we conducted a linear regression to test whether mean (ΔLTL)^2^ is different across relationship types (including parent-offspring pairs).

Using the meta-analysis association estimate, we used the approach described in Visscher et al., ^40^ to estimate the narrow sense heritability (h^2^) attributed to the proportion of shared telomeres IBD (T_shared_). The regression parameter (β) for the association between T_shared_ proportion and (ΔLTL)^2^ is equal to twice the additive genetic variance, and h^2^ is equal to β divided by the twice the total phenotypic variance, multiplied by -1:

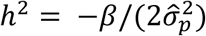

We used this equation to convert the meta-analysis regression parameter β and its upper and lower bound to the h^2^ scale.

In order to determine if LTL reflects TL in germ cells (as expected under the causal diagram in **Figure 1**), used tissue-specific TL data from the Genotype-Tissue Expression (GTEx) project ^42^ to examine the correlation between TL in whole blood and TL in testicular tissue. Tissue collection from post-mortem tissue donors and DNA extraction has been described previously ^43^. TL was measured for GTEx DNA samples using the Luminex technology described above.

## RESULTS

Characteristics of the 5,069 Bangladeshi participants included in our regression analyses are described in **Table 1**. Age was inversely associated with LTL across each cohort (HEALS: r = −0.20, *P* < 10^−16^, BEST: r = −0.23, *P* < 10^−16^, and ACE: r = −0.28, *P* < 10^−16^), as expected. The types of relative pairs observed in our sample are described in **Figure 3**. For each relative pair type, LTL showed significant correlation between relative pairs, with stronger correlation generally observed for closer relative pair types (e.g., siblings r = 0.22, *P* = 3×10^−8^) compared to more distant relatives (i.e., cousins r=0.07, *P* = 0.04) (**Table 2**). Similarly, the mean of our primary outcome of interest, (ΔLTL)^2^, increased with decreasing level of relatedness (**Figure S2**). Univariate regression analyses of (ΔLTL)^2^ on relationship type showed a similar trend, with a larger effect of relationship type for increasing degree of relatedness (**Table S3**).

**Table 1.** Characteristics of Bangladeshi cohorts by type of telomere length measurement: qPCR and the Luminex method (n=5,075)

|  |  | qPCR | Luminex method |  |
| --- | --- | --- | --- | --- |
|  |  | HEALS<br>(2,203) | BEST<br>(1,825) | ACE<br>(1,047) |
| Characteristic |  |  |  |  |
| **Sex** | Male | 1,060 (48.1%) | 996 (54.6%) | 396 (37.8%) |
|  | Female | 1,143 (51.9%) | 829 (45.4%) | 651 (62.2%) |
| **Age (years)** | Mean (SD) | 38.3 (10.6) | 43.7 (10.6) | 36.8 (10.7) |
| **BMI (kg/m<sup>2</sup>)** | <18.5 | 863 (39.6%) | 698 (38.3%) | 431 (41.4%) |
|  | 18.5-24.9 | 1,173 (53.8%) | 963 (52.8%) | 549 (52.7%) |
|  | >25 | 145 (6.7%) | 162 (8.9%) | 62 (5.9%) |
| **Smoking** | Never | 1,298 (58.9%) | 1,110 (60.8%) | 735 (70.2%) |
|  | Former | 176 (8%) | 198 (10.9%) | 58 (5.5%) |
|  | Current | 729 (33.1%) | 517 (28.3%) | 254 (24.3%) |
| **Telomere length** | Mean T/S or TQI<sup>a</sup> | 0.765 (0.17) | 0.672 (0.12) | 0.679 (0.12) |
| **Adjusted TL** | Mean of standardized residuals<sup>b</sup> | 0.02 (0.99) | 0.00 (0.96) | 0.00 (1.06) |

<sup>a</sup>These are the raw T/S ratios (for qPCR) and TQI (for the luminex method) measures, prior to mixed effects modeling.
<sup>b</sup> Standardized residuals from mixed-effects models used to generate difference in telomere length (TL<sub>diff</sub>) variable

**Table 2.** Telomere length correlations for relative pairs by relationship type.

| **Relationship** | **n** | **r<sup>a</sup>** | **P** |
| --- | --- | --- | --- |
| Parent-Offspring | 297 | 0.15 | 0.01 |
| Sibs | 594 | 0.22 | 3.0x10<sup>-8</sup> |
| Avuncular | 606 | 0.09 | 0.02 |
| Cousins | 946 | 0.07 | 0.04 |
| Distant | 5,121 | 0.03 | 0.01 |

<sup>a</sup> Correlations are based on age/sex-adjusted standardized residuals from mixed-effects models.

The total number of chromosome ends shared IBD by relative pairs at each chromosome end (described in **Table S2**) showed evidence of a negative correlation with the cM distance at each chromosome end encompassed by the 180 independent SNPs used to measure IBD sharing (**Figure S3**). This is expected, as chromosome ends with SNP intervals that span a large genetic distance will have more recombination events occurring in that interval than ends with SNPs spanning a shorter genetic distance, resulting in less frequent observation of IBD sharing for the entire segment. The proportion of chromosome ends shared IBD for a relative pair, a proxy for shared telomeres (T_shared_), showed a strong positive association with the estimated pair-wise genetic relatedness (ϕ, estimated using genome-wide SNPs) for each type of relative pair (**Figure 4**). This is also expected, as pairs sharing more of the total genome would be expected to share more telomeres.

**Figure 4.**
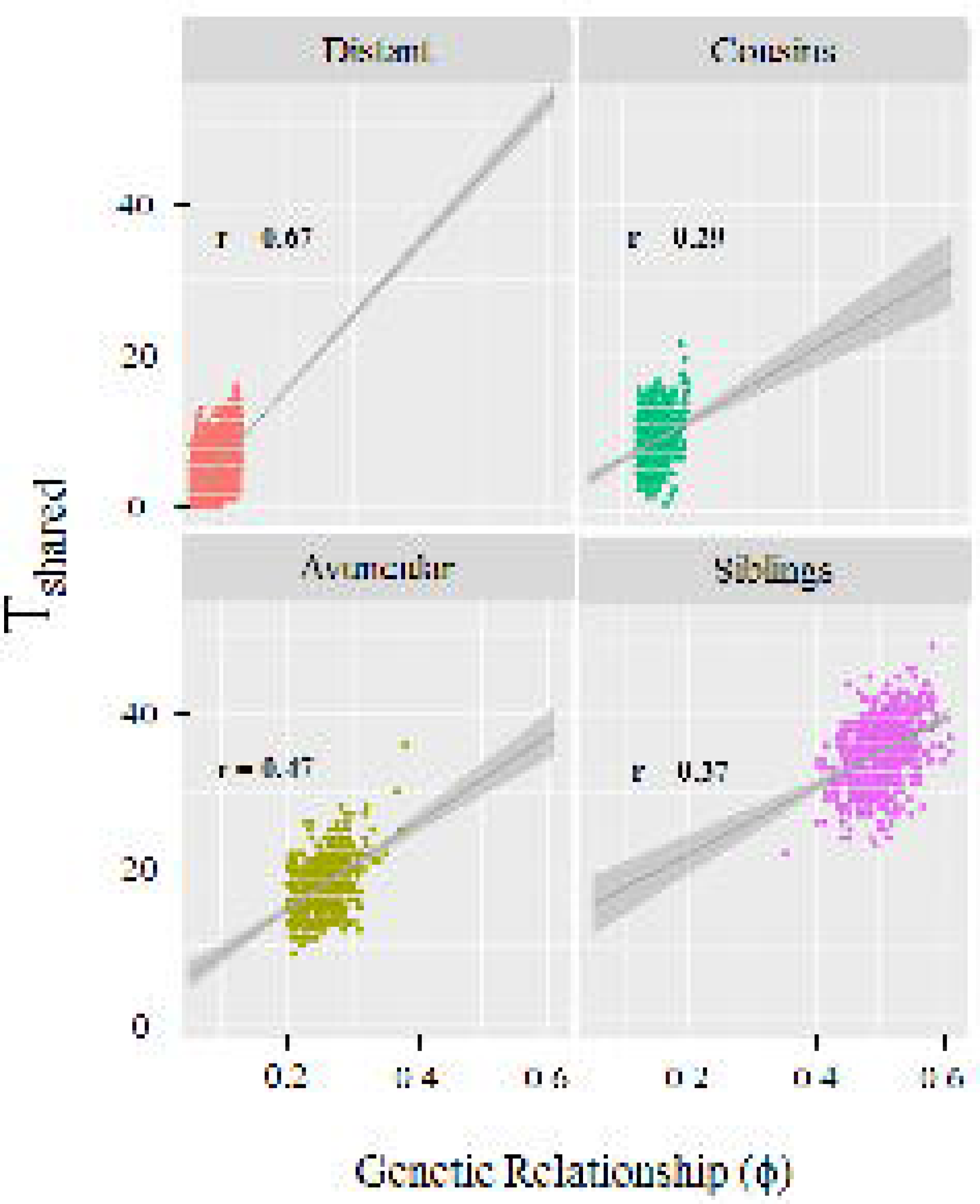
The number of shared telomeres IBD (T_shared_) is strongly correlated with genetic relationship (ϕ) for all four relative pair types. Scatterplots show T_shared_ (y-axis) increases with genetic relatedness (ϕ) (x-axis). Linear fitted lines with 95% confidence intervals and correlation coefficient (r) are included on each panel.

Among each relative pair type, our estimate of the proportion of telomeres shared IBD (the T_shared_ proportion) showed a non-significant inverse association with (ΔLTL)^2^ (**Figure 5**). In contrast, the associations between genome-wide IBD sharing (ϕ) and (ΔLTL)^2^ were generally weaker and did not show a consistent direction of association (**Table 3**). We meta-analyzed the four groups of relative pairs (7,254 total pairs) using a fixed-effects model (test of heterogeneity Q = 2.41, *P* = 0.49), and the T_shared_ proportion showed a significant inverse association with (ΔLTL)^2^ (β = −2.76, *P* = 0.002) (**Figure 5**), while ϕ did not (β = −1.16, *P* = 0.448) (**Table 3**). Using a different threshold to separate cousin pairs from distant pairs (ϕ ≤ 0.08) or combining them into a single group had minimal impact on this result (**Supplementary Figures 4-5**). Restricting to relative pairs for which there were no duplicate individuals resulted in directionally consistent results for T_shared_ (P=0.07) and ϕ (P=0.49) in the meta-analysis, although power was reduced due to smaller sample size (**Supplementary Table 4**).

**Table 3.**
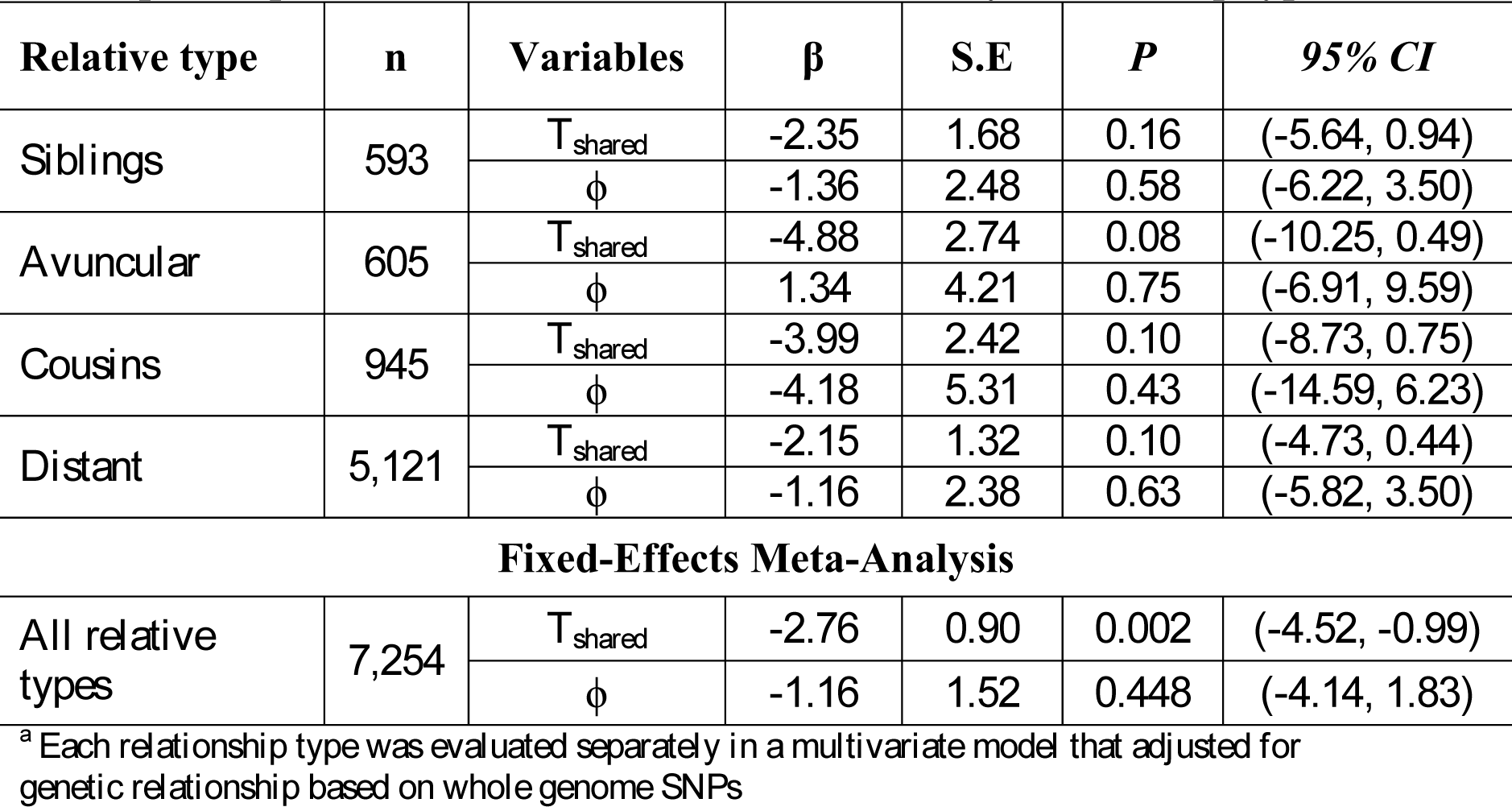
Association of proportion of telomere sharing (T_shared_) and genetic relatedness (ϕ) with squared pairwise difference in LTL ((ΔLTL)^2^) by relationship type^a^

**Figure 5.**
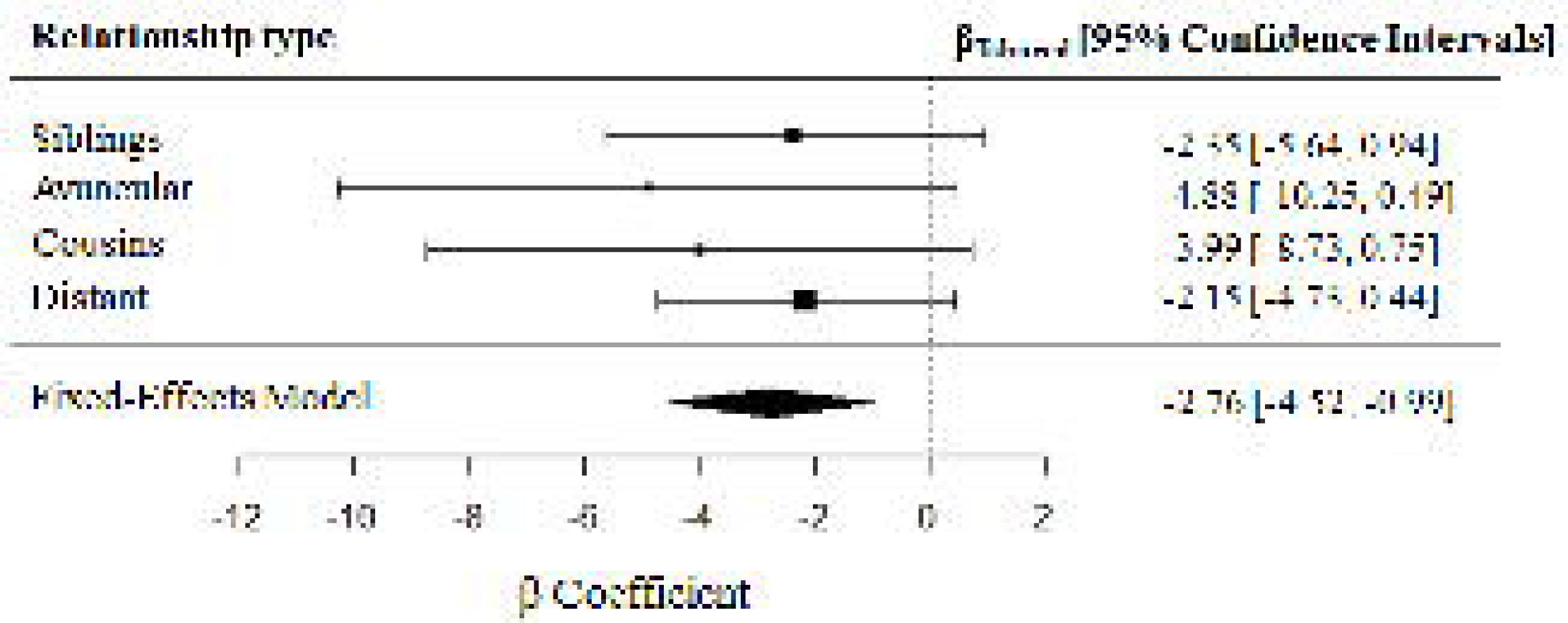
Forest plot of the association between the proportion of telomere shared (T_shared_ proportion) with the squared pairwise difference in LTL ((ΔLTL)^2^), adjusting for genetic relatedness (ϕ). The horizontal bars represent the 95% CI for each association estimate.

Regression analyses using |ΔLTL| as the outcome rather than (ΔLTL)^2^ produced similar results (P=0.0002) (**Table S5, Figure S6**). Scatterplots of the T_shared_ proportion against (ΔLTL)^2^ for each relative pair type are shown in **Supplementary Figure S7-S8**. Together, these analyses indicate that sharing telomeres IBD contributes to pairwise similarity in LTL.

We conducted similar analyses for five continuous control phenotypes to show that the observed association with T_shared_ was specific to pairwise differences in LTL. These phenotypes were height, BMI, mean arterial pressure, systolic blood pressure, and diastolic blood pressure. Difference measures for each phenotype were taken after each phenotype was adjusted for age, sex, cohort, and batch. We did not observe a statistically significant inverse association between the T_shared_ proportion and the pairwise squared difference or the absolute difference for any of the control phenotypes (**Table S6 and S7**).

According to Visscher’s method for h^2^ estimation using genetic data on full siblings ^40^, the narrow-sense h^2^ for the fixed-effects meta-analysis β coefficient is 1.38 (95% CI [0.49, 2.26]). This estimate is above the theoretical maximum for h^2^, likely due to the very large confidence bounds on this estimate. The lower bound for our h^2^ estimate is 0.49, suggesting that at least 49% of the variation in LTL can be explained by sharing telomeres sequence IBD with close relatives. However, variation due to age and other covariates was removed from our TL measures prior to analysis, so the true lower bound is likely lower than 49%.

In order to provide evidence that TL in germ cells and in blood cells reflect TL in the embryonic cells from which both cell types descend (**Figure 1**), we used data on 150 tissue donors from the GTEx project to examine the within-person correlation between TL in whole blood and TL in testicular tissue, which primarily consists of spermatagonia, spermatocytes, or spermatids (~80% of cells). Whole blood TL was positively correlated with TL measured in testicular tissue (r = 0.19, P = 0.02) (**Figure S9**). This observation supports the hypothesized causal relationships in **Figure 1** and the interpretation of our results as trans-generational, direct inheritance of TL.

## DISCUSSION

In this study of LTL measured in a genotyped cohort with substantial relatedness, we show that the extent of sharing chromosome ends IBD, a proxy for sharing telomere sequence IBD, is associated with similarity in LTL (smaller value of (ΔLTL)^2^) measured in a DNA sample. Our results provide evidence that direct transmission of telomeres from parent to offspring accounts for a substantial proportion of LTL h^2^. This implies that variation in TL in parental germ cells impacts TL in offspring embryonic cells (and adult cells) despite telomere reprogramming occurring during embryonic development. The novel approach we describe and apply in this work represents the first attempt to estimate the h^2^ in LTL due to direct transmission of telomeres in a human population.

The association between telomere sharing (T_shared_) and pairwise LTL difference was estimated among sibling, avuncular, cousin, and distant relative pairs. Each of these analyses showed a consistent inverse association, and meta-analysis of all relative pair types produced a statistically significant inverse association. This association was consistently larger than the association between overall genetic relatedness (genome-wide) and LTL difference, suggesting the contribution of telomere sharing to LTL similarity between close relatives is larger than the contribution of sharing inherited variation in non-telomeric regions (e.g., SNPs that influence telomere maintenance).

Genome-wide association (GWA) studies of TL have demonstrated that common genetic variants contribute impact LTL ^34;44-52^. However, the SNPs identified to date account for a <4% of the variation in LTL^17^. Heritability estimates of TL typically range between 34-50% in family studies and have been as high as 82% in twin studies ^13-16^. SNP-based heritability (i.e., variation in TL explained by common SNPs) has been estimated to be ~28% ^53^. These findings suggest additional sources of h^2^, such as the direct transmission of telomeres investigated in this work.

Several prior studies provide support for the hypothesis that direct transmission of telomeres impacts TL. First, there are multiple studies that report a link between paternal age and offspring LTL ^23;24;54-56^. In light of the evidence that telomeres in sperm lengthen as men age, these results suggest that acquired variation in paternal germ cell TL impacts offspring TL in somatic tissues. Additionally, Eisenberg and colleagues ^23^ found the paternal age effect is cumulative across at least two generations, with grandfather’s paternal age (at father’s birth) significantly predicting grandchildren’s TL independent of father’s age at reproduction ^23^. Stindl ^57^ offers a different explanation for the paternal age effect that does not involve age-related telomere lengthening in sperm. Stindl proposes that the association between paternal age and offspring TL is the result of confounding by birth cohort, claiming that TL in the female germline decreases with age and oocytes used first (at younger ages) have longer TL. Consequently, older men have longer sperm TL because they are part of a generation born to younger mothers ^57^. Both explanations for the paternal age effect support the hypothesis that parental transmission of telomeres contributes to TL h^2^.

The work of Aubert *et al*. ^27^, Collopy *et al*. ^28^ and Chiang *et al*. ^29^ also suggests an effect of direct transmission of parental telomeres in humans. These studies report that parents with mutations in *TERT* or *TERC* that render low telomerase activity and shorter telomeres pass their short telomeres to offspring who do not carry the telomerase mutation ^27; 28^. However, these studies were conducted in relatively small samples of parents with telomerase deficiency and their offspring. In addition, these findings were complicated by differences in TL among individuals and the admixing of affected or unaffected paternal and maternal telomeres. ^29^ Thus, these papers provide consistent evidence of the impact of direct transmission on TL, but the implications of these results beyond telomerase-deficient individuals and their families are unclear.

It is worth noting that Chiang et al. ^29^ also reached a similar conclusion based on a telomerase deficient (*Tert*^*+/-*^) mouse model. These mice had critically short and dysfunctional telomeres, whereas their genotypically normal *Tert*^+/+^ offspring had functional telomeres that were protected from progressive shortening but had similar lengths to *Tert*^*+/-*^parents.

To our knowledge, this is the first study to attempt to assess the relationship between IBD sharing of telomeres and similarity in LTL. We focused our analysis on specific types of close relative pairs (siblings, avuncular pairs, cousins, and distant relatives) for several reasons. First, the strength of the presumed relationship between telomere sharing and LTL similarity may weaken as relatives become more distant due to “noise” in TL introduced by telomere reprogramming and other factors affecting TL in germ cells. Second, when examining all types of relative pairs in a single analysis, the correlation between genome-wide relatedness and telomere sharing is very strong (e.g. r=0.97, **Supplementary Figure S10**), and regression models are unable to estimate both parameters; the correlation is weaker for specific relative pair types (**Figure 4**), enabling estimation of both parameters. Also, very distantly-related pairs have an increased number of recombination events occurring since their most recent common ancestor, which makes inferring IBD relationships at chromosome ends more difficult due to more recombination events occurring within the span of SNPs used to estimate IBD. Lastly, traditional SNP-based h^2^ analyses that leverage very distant relationships are not ideal for detecting the effects of telomere sharing on TL similarity because we have no evidence showing that the last SNP measured on each chromosome will be in strong LD with variation in the TL on that chromosome (given the gaps can be hundreds of kb).

Our study utilized two different methods for TL measurement, qPCR and a Luminex-based method. Although we have demonstrated that both methods produce measures that are strongly correlated with Southern blot measures (the gold standard for TL measurement using extracted DNA) ^37; 38^, using multiple methods can introduce heterogeneity into our TL measurement. Therefore, we removed experimental variation from our TL measures through mixed-effects modeling, and standardized TL distributions across all batches and methods to harmonize these measures. In addition, the measures used in this study are a relative measure of average TL (i.e., telomere abundance) rather than an absolute measure of average TL.

Our study has limited power. Assuming there is attenuation in the effect of T_shared_ on (ΔLTL)^2^ with decreasing degree of relatedness, we may be underpowered to detect this difference. Our ability to estimate heritability through the method presented by Visscher *et al.* ^40^ is limited by our small sample size. Visscher *et al*. show that at least 10,000 sibling pairs are required to accurately estimate heritability using the Haseman-Elston-like regression approach. As expected, our h^2^ estimate obtained from the meta-analysis β coefficient has a very wide confidence interval, and produced a point estimate greater than 1. Thus, our work essentially provides a lower bound of the narrow-sense h^2^. Regarding generalizability, it is important to note that these results were obtained using data from a rural Bangladeshi population and cross-population differences in LTL h^2^ could exist due to genetic, environmental, and demographic factors.

To better understand the direct transmission of TL from parents to offspring, future studies should analyze larger numbers of relative pairs, which would enable more accurate estimation of h^2^ due to direct transmission. Our method could be applied to UK Biobank data, which includes a substantial number of relative pairs, as well as LTL data in an upcoming release. Larger studies would also allow researchers to determine if the effect of telomere sharing on pairwise TL differences is weaker for more distant relatives. More accurate methods for measuring TL could increase power for these analyses (e.g., Southern blot), and chromosome-specific measures would enable one to study the transmission of specific telomeres and the impact on chromosome-specific TLs, although these approaches are not currently practical for very large cohorts.

In summary, we provide evidence supporting the direct transmission of variation in TL from parent to offspring. This is the first study to attempt to detect and/or estimate the h^2^ in LTL that is due to direct transmission. Together with prior evidence ^18; 23; 25-27; 54-56^, our work implies that any effect on TL in germ cells – be it due to genes, environment, or aging – will impact offspring TL. Such effects could potentially shift TL in a population in response to demographic or environmental changes or amplify the impact of natural selection on variation (i.e., SNPs) affecting telomere maintenance ^58^.These findings are also relevant for complex diseases for which TL has a causal impact on disease risk, as the h^2^ for these traits may be impacted by trans-generationally inherited TL, a component of h^2^ that cannot be explained by SNPs or other variation in non-telomere sequences. Applying our method to larger studies of relative pairs will enable robust estimation of LTL h^2^ attributable to direction transmission.

## SUPPLEMENTAL DATA

Supplemental data (included as a separate pdf file) include seven tables and ten figures.

### ACKNOWLEDGMENTS

This work was supported by the National Institutes of Health [R01 ES020506, U01 HG007601, P42 ES10349, R01 CA107431, R01 CA102484, P30 CA014599]. The authors thank all the men and women who participated in HEALS and BEST and all research staff who contributed to data collection. The Genotype-Tissue Expression (GTEx) Project was supported by the Common Fund of the Office of the Director of the National Institutes of Health, and by NCI, NHGRI, NHLBI, NIDA, NIMH, and NINDS.

## REFERENCES

1. Blackburn, E.H., Epel, E.S., and Lin J. (2015). Human telomere biology: A contributory and interactive factor in aging, disease risks, and protection. Science 350, 1193–1198.

2. Aubert, G., and Lansdorp P.M. (2008). Telomeres and Aging. Physiological Reviews 88, 557–579.

3. Huang, Y., Liang, P., Liu, D., Huang, J., and Songyang Z. (2014). Telomere regulation in pluripotent stem cells. Protein & Cell 5, 194–202.

4. Cawthon, R.M., Smith, K.R., O’Brien, E., Sivatchenko, A., and Kerber, R.A. (2003). Association between telomere length in blood and mortality in people aged 60 years or older. The Lancet 361, 393–395.

5. Haycock, P.C., Heydon, E.E., Kaptoge, S., Butterworth, A.S., Thompson, A., and Willeit, P. (2014). Leucocyte telomere length and risk of cardiovascular disease: systematic review and meta-analysis. BMJ: British Medical Journal 349.

6. Willeit, P., Willeit, J., Brandstätter, A., Ehrlenbach, S., Mayr, A., Gasperi, A., Weger, S., Oberhollenzer, F., Reindl, M., Kronenberg, F., et al (2010). Cellular Aging Reflected by Leukocyte Telomere Length Predicts Advanced Atherosclerosis and Cardiovascular Disease Risk. Arteriosclerosis, Thrombosis, and Vascular Biology 30, 1649–1656.

7. Willeit, P., Willeit, J., Mayr, A., and et al (2010). Telomere length and risk of incident cancer and cancer mortality. JAMA 304, 69–75.

8. Zhang, C., Doherty, J.A., Burgess, S., Hung, R.J., Lindström, S., Kraft, P., Gong, J., Amos, C.I., Sellers, T.A., Monteiro, A.N.A., et al (2015). Genetic determinants of telomere length and risk of common cancers: a Mendelian randomization study. Human molecular genetics 24, 5356–5366.

9. Iles, M.M., Bishop, D.T., Taylor, J.C., Hayward, N.K., Brossard, M., Cust, A.E., Dunning, A.M., Lee, J.E., Moses, E.K., Akslen, L.A., et al (2014). The effect on melanoma risk of genes previously associated with telomere length. Journal of the National Cancer Institute 106.

10. Walsh, K.M., Codd, V., Rice, T., Nelson, C.P., Smirnov, I.V., McCoy, L.S., Hansen, H.M., Elhauge, E., Ojha, J., Francis, S.S., et al (2015). Longer genotypically-estimated leukocyte telomere length is associated with increased adult glioma risk. Oncotarget 6, 42468–42477.

11. Walsh, K.M., Whitehead, T.P., de Smith, A.J., Smirnov, I.V., Park, M., Endicott, A.A., Francis, S.S., Codd, V., Samani, N.J., Metayer, C., et al (2016). Common genetic variants associated with telomere length confer risk for neuroblastoma and other childhood cancers. Carcinogenesis 37, 576–582.

12. Ojha, J., Codd, V., Nelson, C.P., Samani, N.J., Smirnov, I.V., Madsen, N.R., Hansen, H.M., de Smith, A.J., Bracci, P.M., Wiencke, J.K., et al (2016). Genetic Variation Associated with Longer Telomere Length Increases Risk of Chronic Lymphocytic Leukemia. Cancer Epidemiol Biomarkers Prev 25, 1043–1049.

13. Bischoff, C., Graakjaer, J., Petersen, H.C., Jeune, B., Bohr, V.A., Koelvraa, S., and Christensen, K. (2012). Telomere Length Among the Elderly and Oldest-Old. Twin Research and Human Genetics 8, 425–432.

14. Hjelmborg, J.B., Dalgård, C., Möller S., Steenstrup, T., Kimura, M., Christensen, K., Kyvik, K.O., and Aviv, A. (2015). The heritability of leucocyte telomere length dynamics. Journal of Medical Genetics.

15. Honig, L.S., Kang, M.S., Cheng, R., Eckfeldt, J.H., Thyagarajan, B., Leiendecker-Foster, C., Province, M.A., Sanders, J.L., Perls, T., Christensen, K., et al (2015). Heritability of telomere length in a study of long-lived families. Neurobiology of Aging 36, 2785–2790.

16. Broer, L., Codd, V., Nyholt, D.R., Deelen, J., Mangino, M., Willemsen, G., Albrecht, E., Amin, N., Beekman, M., de Geus, E.J.C., et al (2013). Meta-analysis of telomere length in 19 713 subjects reveals high heritability, stronger maternal inheritance and a paternal age effect. European Journal of Human Genetics 21, 1163–1168.

17. Codd, V., Nelson, C.P., Albrecht, E., Mangino, M., Deelen, J., Buxton, J.L., Hottenga, J.J., Fischer, K., Esko, T., Surakka, I., et al (2013). Identification of seven loci affecting mean telomere length and their association with disease. Nat Genet 45, 422–427.

18. De Meyer, T., Vandepitte, K., Denil, S., De Buyzere, M.L., Rietzschel, E.R., and Bekaert, S. (2014). A non-genetic, epigenetic-like mechanism of telomere length inheritance? European Journal of Human Genetics 22, 10–11.

19. Kalmbach, K., Robinson, L.G. Jr., Wang, F., Liu, L., and Keefe, D. (2014). Telomere length reprogramming in embryos and stem cells. Biomed Res Int 2014, 925121.

20. Kalmbach, K.H., Antunes, D.M.F., Dracxler, R.C., Knier, T.W., Seth-Smith, M.L., Wang, F., Liu, L., and Keefe, D.L. (2013). Telomeres and human reproduction. Fertility and sterility 99, 10.1016/j.fertnstert.2012.1011.1039.

21. Liu, L., Trimarchi, J.R., Smith, P.J., and Keefe, D.L. (2002). Mitochondrial dysfunction leads to telomere attrition and genomic instability. Aging cell 1, 40–46.

22. Allsopp, R.C., Vaziri, H., Patterson, C., Goldstein, S., Younglai, E.V., Futcher, A.B., Greider, C.W., and Harley, C.B. (1992). Telomere length predicts replicative capacity of human fibroblasts. Proc Natl Acad Sci U S A 89, 10114–10118.

23. Eisenberg, D.T.A., Hayes, M.G., and Kuzawa, C.W. (2012). Delayed paternal age of reproduction in humans is associated with longer telomeres across two generations of descendants. Proceedings of the National Academy of Sciences 109, 10251–10256.

24. Kimura, M., Cherkas, L.F., Kato, B.S., Demissie, S., Hjelmborg, J.B., Brimacombe, M., Cupples, A., Hunkin, J.L., Gardner, J.P., Lu, X., et al (2008). Offspring’s Leukocyte Telomere Length, Paternal Age, and Telomere Elongation in Sperm. PLOS Genetics 4, e37.

25. Prescott, J., Du, M., Wong, J.Y., Han, J., and De Vivo, I. (2012). Paternal age at birth is associated with offspring leukocyte telomere length in the nurses’ health study. Human reproduction (Oxford, England) 27, 3622–3631.

26. Achi, M.V., Ravindranath, N., and Dym, M. (2000). Telomere length in male germ cells is inversely correlated with telomerase activity. Biology of reproduction 63, 591–598.

27. Aubert, G., Baerlocher, G.M., Vulto, I., Poon, S.S., and Lansdorp, P.M. (2012). Collapse of Telomere Homeostasis in Hematopoietic Cells Caused by Heterozygous Mutations in Telomerase Genes. PLOS Genetics 8, e1002696.

28. Collopy, L.C., Walne, A.J., Cardoso, S., de la Fuente, J., Mohamed, M., Toriello, H., Tamary, H., Ling, A.J.Y.V., Lloyd, T., Kassam, R., et al (2015). Triallelic and epigenetic-like inheritance in human disorders of telomerase. Blood 126, 176.

29. Chiang, Y.J., Calado, R.T., Hathcock, K.S., Lansdorp, P.M., Young, N.S., and Hodes, R.J. (2010). Telomere length is inherited with resetting of the telomere set-point. Proceedings of the National Academy of Sciences 107, 10148–10153.

30. Ahsan, H., Chen, Y., Parvez, F., Argos, M., Hussain, A.I., Momotaj, H., Levy, D., van Geen, A., Howe, G., and Graziano, J. (2005). Health Effects of Arsenic Longitudinal Study (HEALS): Description of a multidisciplinary epidemiologic investigation. J Expos Sci Environ Epidemiol 16, 191–205.

31. Argos, M., Rahman, M., Parvez, F., Dignam, J., Islam, T., Quasem, I., Hore, S.K., Haider, A.T., Hossain, Z., Patwary, T.I., et al (2013). Baseline Comorbidities in a Skin Cancer Prevention Trial in Bangladesh. European journal of clinical investigation 43, 579–588.

32. Pierce, B.L., Kibriya, M.G., Tong, L., Jasmine, F., Argos, M., Roy, S., Paul-Brutus, R., Rahaman, R., Rakibuz-Zaman, M., Parvez, F., et al (2012). Genome-Wide Association Study Identifies Chromosome 10q24.32 Variants Associated with Arsenic Metabolism and Toxicity Phenotypes in Bangladesh. PLoS Genetics 8, e1002522.

33. Pierce, B.L., Tong, L., Argos, M., Gao, J., Jasmine, F., Roy, S., Paul-Brutus, R., Rahaman, R., Rakibuz-Zaman, M., Parvez, F., et al (2013). Arsenic metabolism efficiency has a causal role in arsenic toxicity: Mendelian randomization and gene-environment interaction. International Journal of Epidemiology 42, 1862–1872.

34. Delgado, D.A., Zhang, C., Chen, L.S., Gao, J., Roy, S., Shinkle, J., Sabarinathan, M., Argos, M., Tong, L., Ahmed, A., et al (2017). Genome-wide association study of telomere length among South Asians identifies a second RTEL1 association signal. Journal of Medical Genetics.

35. Ehrlenbach, S., Willeit, P., Kiechl, S., Willeit, J., Reindl, M., Schanda, K., Kronenberg, F., and Brandstätter, A. (2009). Influences on the reduction of relative telomere length over 10 years in the population-based Bruneck Study: introduction of a well-controlled high-throughput assay. International Journal of Epidemiology 38, 1725–1734.

36. Cawthon, R.M. (2002). Telomere measurement by quantitative PCR. Nucleic Acids Research 30, e47–e47.

37. Kibriya, M.G., Jasmine, F., Roy, S., Ahsan, H., and Pierce, B. (2014). Measurement of Telomere length: a new assay using QuantiGene chemistry on a Luminex platform. Cancer epidemiology, biomarkers & prevention: a publication of the American Association for Cancer Research, cosponsored by the American Society of Preventive Oncology 23, 2667–2672.

38. Pierce, B.L., Jasmine, F., Roy, S., Zhang, C., Aviv, A., Hunt, S.C., Ahsan, H., and Kibriya, M.G. (2016). Telomere length measurement by a novel Luminex-based assay: a blinded comparison to Southern blot. International Journal of Molecular Epidemiology and Genetics 7, 18–23.

39. Visscher, Peter M., Macgregor, S., Benyamin, B., Zhu, G., Gordon, S., Medland, S. Hill, William G., Hottenga, J.-J., Willemsen, G., Boomsma, Dorret I., et al (2007). Genome Partitioning of Genetic Variation for Height from 11,214 Sibling Pairs. American Journal of Human Genetics 81, 1104–1110.

40. Visscher, P.M., Medland, S.E., Ferreira, M.A.R., Morley, K.I., Zhu, G., Cornes, B.K., Montgomery, G.W., and Martin, N.G. (2006). Assumption-Free Estimation of Heritability from Genome-Wide Identity-by-Descent Sharing between Full Siblings. PLOS Genetics 2, e41.

41. Hemani, G., Yang, J., Vinkhuyzen, A. Powell, Joseph E., Willemsen, G., Hottenga, J.-J., Abdellaoui, A., Mangino, M., Valdes, Ana M., Medland, Sarah E., et al (2013). Inference of the Genetic Architecture Underlying BMI and Height with the Use of 20,240 Sibling Pairs. American Journal of Human Genetics 93, 865–875.

42. eGTEx Project. (2017). Enhancing GTEx by bridging the gaps between genotype, gene expression, and disease. Nature Genetics 49, 1664.

43. Carithers, L.J., Ardlie, K., Barcus, M., Branton, P.A., Britton, A., Buia, S.A., Compton, C.C., DeLuca, D.S., Peter-Demchok, J., Gelfand, E.T., et al (2015). A Novel Approach to High-Quality Postmortem Tissue Procurement: The GTEx Project. Biopreservation and biobanking 13, 311–319.

44. Codd, V., Mangino, M., van der Harst, P., Braund, P.S., Kaiser, M., Beveridge, A.J., Rafelt, S., Moore, J., Nelson, C., Soranzo, N., et al (2010). Common variants near TERC are associated with mean telomere length. Nat Genet 42, 197–199.

45. Do, S.K., Yoo, S.S., Choi, Y.Y., Choi, J.E., Jeon, H.-S., Lee, W.K., Lee, S.Y., Lee, J., Cha, S.I., Kim, C.H., et al (2015). Replication of the results of genome-wide and candidate gene association studies on telomere length in a Korean population. The Korean Journal of Internal Medicine 30, 719–726.

46. Levy, D., Neuhausen, S.L., Hunt, S.C., Kimura, M., Hwang, S.-J., Chen, W., Bis, J.C., Fitzpatrick, A.L., Smith, E., Johnson, A.D., et al (2010). Genome-wide association identifies OBFC1 as a locus involved in human leukocyte telomere biology. Proceedings of the National Academy of Sciences of the United States of America 107, 9293–9298.

47. Mangino, M., Christiansen, L., Stone, R., Hunt, S.C., Horvath, K., Eisenberg, D.T.A., Kimura, M., Petersen, I., Kark, J.D., Herbig, U., et al (2015). DCAF4, a novel gene associated with leucocyte telomere length. Journal of Medical Genetics 52, 157–162.

48. Mangino, M., Hwang, S.-J., Spector, T.D., Hunt, S.C., Kimura, M., Fitzpatrick, A.L., Christiansen, L., Petersen, I., Elbers, C.C., Harris, T., et al (2012). Genome-wide meta-analysis points to CTC1 and ZNF676 as genes regulating telomere homeostasis in humans. Human molecular genetics 21, 5385–5394.

49. Pooley, K.A., Bojesen, S.E., Weischer, M., Nielsen, S.F., Thompson, D., Amin Al Olama, A., Michailidou, K., Tyrer, J.P., Benlloch, S., Brown, J., et al (2013). A genome-wide association scan (GWAS) for mean telomere length within the COGS project: identified loci show little association with hormone-related cancer risk. Human molecular genetics 22, 5056–5064.

50. Prescott, J., Kraft, P., Chasman, D.I., Savage, S.A., Mirabello, L., Berndt, S.I., Weissfeld, J.L., Han, J., Hayes, R.B., Chanock, S.J., et al (2011). Genome-wide association study of relative telomere length. PloS one 6, e19635.

51. Shen, Q., Zhang, Z., Yu, L., Cao, L., Zhou, D., Kan, M., Li, B., Zhang, D., He, L., and Liu, Y. (2011). Common variants near TERC are associated with leukocyte telomere length in the Chinese Han population. Eur J Hum Genet 19, 721–723.

52. Soerensen, M., Thinggaard, M., Nygaard, M., Dato, S., Tan, Q., Hjelmborg, J., Andersen-Ranberg, K., Stevnsner, T., Bohr, V.A., Kimura, M., et al (2012). Genetic variation in TERT and TERC and human leukocyte telomere length and longevity: a cross sectional and longitudinal analysis. Aging cell 11, 223–227.

53. Faul, J.D., Mitchell, C.M., Smith, J.A., and Zhao, W. (2016). Estimating Telomere Length Heritability in an Unrelated Sample of Adults: Is Heritability of Telomere Length Modified by Life Course Socioeconomic Status? Biodemography and social biology 62, 73–86.

54. Unryn, B.M., Cook, L.S., and Riabowol, K.T. (2005). Paternal age is positively linked to telomere length of children. Aging cell 4, 97–101.

55. Broer, L., Codd, V., Nyholt, D.R., Deelen, J., Mangino, M., Willemsen, G., Albrecht, E., Amin, N., Beekman, M., de Geus, E.J.C., et al (2013). Meta-analysis of telomere length in 19[thinsp]713 subjects reveals high heritability, stronger maternal inheritance and a paternal age effect. Eur J Hum Genet 21, 1163–1168.

56. De Meyer, T., Rietzschel, E.R., De Buyzere, M.L., De Bacquer, D., Van Criekinge, W., De Backer, G.G., Gillebert, T.C., Van Oostveldt, P., and Bekaert, S. (2007). Paternal age at birth is an important determinant of offspring telomere length. Human molecular genetics 16, 3097–3102.

57. Stindl, R. (2016). The paradox of longer sperm telomeres in older men’s testes: a birth-cohort effect caused by transgenerational telomere erosion in the female germline. Molecular Cytogenetics 9, 12.

58. Hansen, M.E., Hunt, S.C., Stone, R.C., Horvath, K., Herbig, U., Ranciaro, A., Hirbo, J., Beggs, W., Reiner, A.P., Wilson, J.G., et al (2016). Shorter telomere length in Europeans than in Africans due to polygenetic adaptation. Human molecular genetics.

